# Bacterial Spores as a Scalable, Modular Platform for the Production of Amyloids for Materials

**DOI:** 10.64898/2026.03.19.712379

**Authors:** Carina Dietz, Maren Kvilten, Simona Sebastiano, Cécile Formosa-Dague, Alexander Unger, Dieter Spiehl, Andreas Blaeser, Mikael Lindgren, Magnus Philipp, Johannes Kabisch

## Abstract

We present a proof-of-concept platform in which amyloids are displayed on the surface of engineered *Bacillus subtilis* spores for bioengineered materials. Amyloids possess high tensile strength, elasticity, and tunable assembly, but their use is limited by inaccessible native sources and low-yield or toxic heterologous expression. Here, spores were engineered to display the native amyloid TasA and Humboldt squid suckerins 9 and 10 as fusions to the spore coat protein CotY. Amyloid production and fibril formation were confirmed by Western blot and X-34 staining, and quantitative analysis indicated mg/L-level yields. Atomic force microscopy revealed altered stiffness and surface ultrastructure, and incorporation of amyloid-displaying spores into resin-based 3D printing modified tensile strength. These findings highlight spore-based amyloid display as a scalable, modular platform for materials applications, leveraging established industrial spore production.

## 1. Introduction

The soil bacterium *Bacillus subtilis* is a well described model organism and industrial workhorse in biotechnology (Stülke et al., 2023). Next to its capability to secrete industrially relevant enzymes (Rajagopalan and Krishnan, 2008; Wei et al., 2012), it is also known as an industrially relevant producer of secondary metabolites (Kaspar et al., 2019; Schwechheimer et al., 2016). *Bacillus subtilis* has been described to produce endospores as early as the 19^th^ century (Cohn, 1877).

**Fig. 1:**
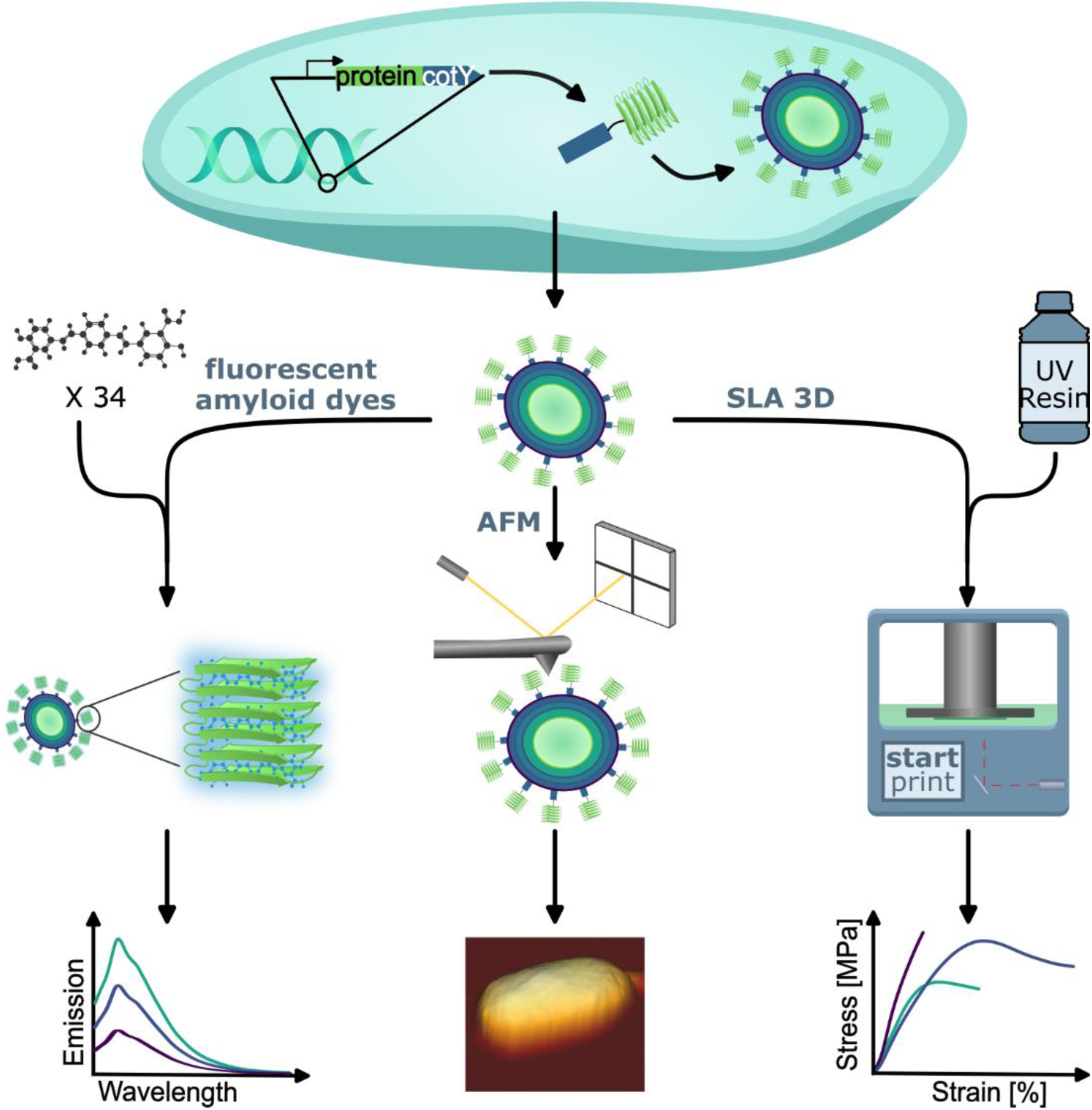
Graphical abstract: Genetic makeup of spore-display constructs used in this study. A gene encoding for a protein of interest is genetically fused either to the future N- or C-terminus of the spore-coat protein encoding gene *cotY*. Gene expression is regulated via the sporulation dependent *cotYZ*-promoter, ensuring protein production in the late stage of sporulation (spore-coat formation). The genetic information is encoded on the high-copy template vector pCascade (Karava et al., 2019). After separation of mother-cell and forespore the spore coat is formed. The coat protein CotY and its designated fusion protein are integrated into the spore coat from the cytosol of the mother cell. After sporulation induced lysis, amyloids on the surface are tested for properties.

Sporulation is a complex process that involves asymmetric cell division, resulting in the formation of the forespore and mother cell. Subsequently, the spore coat is assembled around the forespore, resulting in the formation of a mature endospore housing a copy of the cells’ chromosome. Finally, the mother cell lyses releasing the endospore into the environment (Tan and Ramamurthi, 2014). Intended as a survival form in hostile or nutrient starved environments, bacterial spores are among the hardiest forms of life on earth. They have been shown to be resistant to temperature, UV radiation, dehydration, and oxidative damage (Nicholson et al., 2000; Setlow, 2016, 1995). Furthermore, bacterial spores have been documented to be resistant to predation by other microorganisms (Klobutcher et al., 2006; Laaberki and Dworkin, 2008) as well as toxic chemicals (Milhaud and Balassa, 1973) and display extreme longevity demonstrated by the recovery of viable spores from environmental samples as old as 10.000 years (Potts, 1994).

This hardiness can largely be attributed to the spore coat. The *B. subtilis* spore coat has a width slightly below 200 nm and is made up of four layers: the basement layer, the inner coat, outer coat and crust. Assembly of the spore coat encompasses very complex structural mechanisms in bacteria and involves 2 % of the entire genome and 15 % of sporulation genes with tight spatiotemporal regulation of the sporulation process in both compartments (Eijlander et al., 2014; Errington and Illing, 1992; LaBell et al., 1987,(Driks and Eichenberger, 2016).

Display of a heterologous protein on the spore surface was first demonstrated using Gfp, after which Isticato et al. demonstrated the display of the C-terminal fragment of the tetanus toxin using CotB on the surface of *B. subtilis* spores (Isticato et al., 2001; Webb et al., 1995). Based on this research, several spore coat and crust proteins have been utilized to display proteins (Guoyan et al., 2019).

Spore surface display combines the advantages of classic protein immobilization with a robust system and simple purification via centrifugation, thus avoiding costly immobilization steps of purified proteins on chemically treated surfaces or the difficult handling of other cell- or phage-based immobilization systems (Chen et al., 2017). Furthermore, it brings with it unique advantages lent by the protective nature of the spore coat like the display and usage of enzymes from extremophiles as evident by the display of a thermostable Esterase-CotB fusion protein that retained stable activity at 70 °C (Chen et al., 2015). Moreover, it has been shown that factors like thermo- and pH stability of proteins displayed on spores are enhanced (He et al., 2025; Ullah et al., 2024). Additionally, increased stability and easy recovery of spores allow for recycling of displayed proteins (Qu et al., 2014; Spannenkrebs et al., 2025).

While spore surface display is a well-established method to produce enzymes and medically relevant proteins, the display of amyloids has so far not been demonstrated. The term “amyloid” describes not a class of proteins but rather a defined structural conformation. Proteins in amyloid conformation display an unbranched fibrillar structure formed of β-sheets that are arranged parallel to the fibril axis (Pauling and Corey, 1951; Sunde et al., 1997). The best studied examples are involved in human amyloidosis, where fibril formation into plaques disrupts normal organ function (Chiti and Dobson, 2017). Well studied amyloids include amyloid-β (Aβ)40/42 associated with Alzheimer’s disease and α-synuclein associated with Parkinson’s disease (Murphy and LeVine, 2010; Stefanis, 2012).

However, despite their role in detrimental human disease, amyloids and amyloid-like proteins have functions in a variety of other organisms, ranging from bacteria to arthropods and mollusks. These include structure giving proteins in biofilms and proteins involved in bacterial adhesion, such as the *E. coli* Curli amyloids, FapC from *Pseudomonas fluorescens* and TasA from *Bacillus subtilis* (Branda et al., 2006; Chapman et al., 2002; Dueholm et al., 2010). Compared to amyloid formation in amyloidosis, which often occurs at random due to misfolding, bacterial amyloid formation is a regulated process involving chaperones and nucleators. In *E. coli,* the formation of Curli fibrils includes at least six proteins (Chapman et al., 2002; Evans et al., 2015; Hammar et al., 1995; Hammer et al., 2007; Nenninger et al., 2011). In *B. subtilis,* TasA formation in the biofilm is facilitated by three proteins, TasA itself, the nucleation factor and chaperone TapA as well as signal peptidase SipW (Bamford et al., 2024; Roske et al., 2023; Terra et al., 2012).

The most interesting class of amyloid-like proteins from a material perspective are load-bearing proteins such as Spidroins that make up spider silk and Suckerins that form the ring teeth of squid. While the fibrils formed by these proteins do not exactly match amyloid fibrils, they form a similar cross-β-structure and have been shown to bind amyloid specific dyes (Hiew et al., 2016; Slotta et al., 2007). Spider silk fibers have been shown to be up to five times stronger than steel and three times tougher than Kevlar (Gosline et al., 1999). However, long, repetitive DNA sequences, formation of inclusion bodies, overall low yield, and reduced tensile strength of synthetic fibers compared to native fibers pose a challenge for heterologous production (Ramezaniaghdam et al., 2022). Squid ring tooth Suckerins (SRT) are less well researched than Spidroins but show highly interesting properties. SRTs have been shown to be very similar in structure to silk-like fibers showing high mechanical strength (Ding et al., 2014). Additionally, naturally extracted SRTs can be solved in watery solution by heating and molded like thermoplastics, which allows for the formation of soft or hard materials based on humidity and water content (Latza et al., 2015; Rieu et al., 2016). Furthermore, SRT solutions in acetic acid can be molded into flexible biofilms or processed into stiff gels and films by ruthenium crosslinking (Ding et al., 2015). Recently, SRTs have been integrated to form chitosan/SRT hydrogels via genipin cross-linking, demonstrating the potential of SRTs in multi-polymer applications (Chen et al., 2025). This broad range of properties can be accredited to their unique structure. Most SRTs have a repetitive sequence of flexible Glycine rich and stiff Alanine, Histidine, Threonine, Valine and Serine rich repeats, although it is to note that some SRT variants display a much higher content of flexible or stiff regions (Guerette et al., 2014). These interchanging flexible and stiff repeats form nano-confined, β-sheet reinforced supramolecular networks. Similarly to Spidroins however, large scale use of SRTs is still held back by difficulties in recombinant production and processing.

Analysis of amyloid formation is often carried out through FT-IR, Raman or circular dichroism spectroscopy, nuclear magnetic resonance (NMR) or cryo electron microscopy (cryo-EM) both in the fields of basic amyloid research and synthetic biology (Asakura et al., 2022; Böhning et al., 2022; Ridgley et al., 2013). Another regularly employed method to assess not only structure but material properties, is atomic force microscopy (AFM) (Adamcik and Mezzenga, 2012). A more accessible alternative can be found with amyloid specific dyes. Congo-Red and Thioflavin-T intercalate with amyloid fibers, resulting in a spectral shift in absorbance or fluorescence, respectively (Khurana et al., 2005; Puchtler et al., 1962). However, Congo-red and Thioflavin-T both have drawbacks, as they have been shown to not be entirely specific to amyloid fibers (Krebs et al., 2005; Yakupova et al., 2019). These drawbacks have led to the development of an array of new dyes ranging from X-34, a derivative of Congo-Red that bind amyloids in both fibril and fiber state, to a toolbox of dyes based on pentameric thiophene scaffolds and pH resistant dyes such as BTD-17 and BTD-21 (Åslund et al., 2009; Styren, et al., 2000; Yuzu et al., 2020; Zhang et al., 2019). Given the current state of the art and methodological possibilities at hand, we decided to develop a new technique for amyloid production, characterization, and potential application. In this study, we set out to develop the production of amyloid-like proteins in bacteria by displaying them on the surface of *B. subtilis* spores, combining the toughness and longevity of bacterial spores with the excellent material properties of amyloid-like proteins with a simple purification procedure. The resulting amyloid displaying spores were characterized and used as an additive to resins in stereolithography (SLA) 3-D printing.

## 2. Results

### 2.1 Amyloid proteins are present on the spore-surface as oligomers

As a first step towards display of amyloid proteins, we generated strains harboring genetic constructs with C- and N-terminal fusions of the *D. gigas* Suckerins (SRT) 9 (uniprot: **A0A075LXZ4)**, 10 (uniprot: **A0A075LY72)** and native amyloid-like biofilm scaffolding protein TasA (uniprot: **P54507)** to the native spore-coat protein CotY. As a control, the enzyme sucrose phosphorylase (SucP, uniprot: **A0ZZH6**) from the bacterium *Bifidobacterium adolescentis* was chosen. The native *Bacillus subtilis* TasA served as a proof of principle, while we chose SRTs 9 and 10 from *Dosidicus gigas* (great Humboldt squid) as representative proteins for the different types of SRT proteins. SRT9 is a Glycine-rich, largely amorphous protein and the partially crystalline SRT10 shows a tandem-repeat structure of interchanging Gly-rich and Ala/His/Thr/Val/Ser-rich repeats flanked by Proline. Plasmids were transformed into germination deficient strains harboring deletions of genes *sleB* and *cwlD* lowering germination rates to 0.001 % (Karava et al., 2019), ensuring high stability of protein bearing spores. To avoid false positive results based on the native production of amyloid-like protein TasA, the *tasA* operon was deleted in display strains. However, this resulted in a reduction in spore yield and loss of protein display in strains displaying TasA on the spore surface (see Fig. 2 and supplemental Fig. 1). To avoid this, genes encoding for auxiliary proteins TapA, a molecular chaperone assisting in TasA filament formation (Roske et al., 2023) and SipW, a putative signal peptidase (Terra et al., 2012) were reconstituted by reintroduction into the *tasA* locus under control of the stationary phase promoters *pylb* and *p2* respectively (Feavers et al., 1988), restoring strain fitness and spore yield (see Fig. 2 and supplemental Fig. 1).

**Fig 2:**
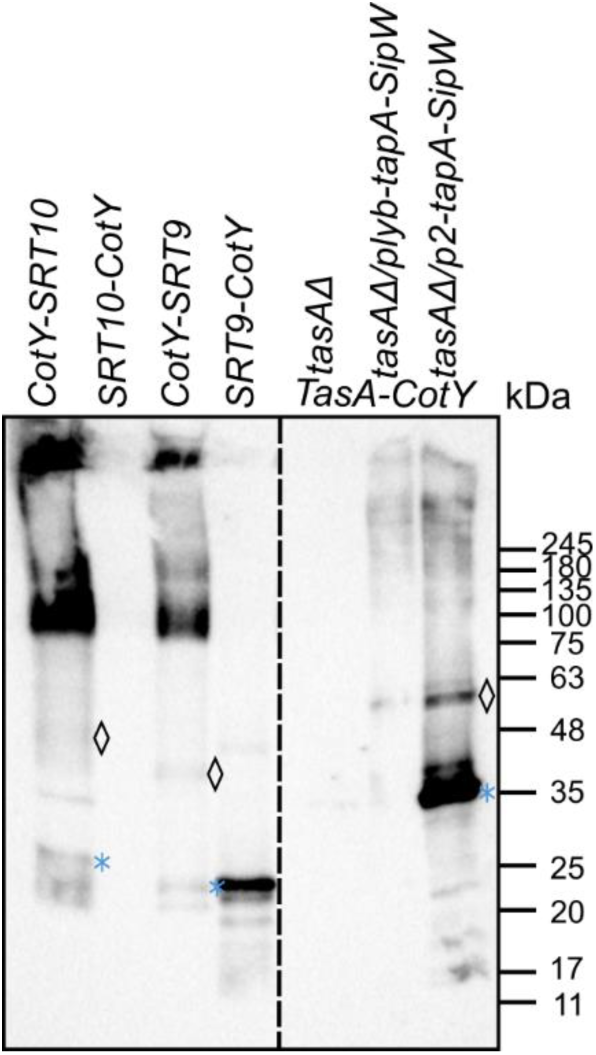
Western blot of the proteins removed from the spore surface. 5 µl samples were separated by SDS-PAGE and subjected to Western blot analysis. Text on top denotes the strain the sample was taken from. Symbols to the left of signals indicate expected protein monomer (asterisk) and CotY-fusion protein (diamond) sizes. CotY-X denotes N-terminal fusion while X-CotY denotes a C-terminal fusion. Protein ladder sizes are shown on the right. The dashed line indicates the position where the image was cropped for clarity. One representative replicate is shown.

To confirm the presence of the target proteins on the spore surface, we generated epitope-tagged versions of these constructs. Spores of these strains were treated with heat, detergents, and bead-beating to strip the spore-coat. Supernatant samples of stripped spores were subjected to SDS-PAGE and Western blot analysis. Western blot analysis revealed a signal in TasA-CotY at roughly 35 kDa corresponding to the expected size of TasA (32 kDa). In addition, a signal was observed at 55 kDa corresponding to the TasA-CotY fusion protein. Additional bands were detected in the range of 100 – 245 kDa likely indicating oligomeric forms of the protein. Notably, no signal was obtained when the native *tasA* operon was deleted. Reconstitution of the operon via *pylb* resulted in moderate signals, while putting the operon under control of *p2* resulted in a strong signal for TasA. No signal was obtained for the N-terminal fusion CotY-TasA, and the strain was subsequently excluded from further experiments. Signals were detected at 48 kDa for SRT10 and above 35 kDa for SRT9 corresponding to the respective sizes of the CotY-fusions, while a signal above 25 kDa likely indicates the 27 kDa SRT10 monomer and a signal above 20 kDa the 22 kDa SRT9 monomer. Both N-terminal SRT proteins showed a higher degree of oligomerization than TasA with strong signals between 75 and 100 kDa, between 135 and 180 kDa and above 245 kDa. Stripping of SRT9-CotY and SRT10-CotYs spores, respectively, yielded only a monomer or no signal at all (Fig. 2).

### 2.1 Displayed proteins form amyloid fibrils

After confirming the production of candidate proteins, we assessed the fold of the target proteins on the spore-surface by the use of amyloid-specific dyes. Staining the spores with Congo-Red was unsuccessful as all types of spores, including controls were stained by the dye (see supplemental Fig. 2). Thioflavin-T was not tested, as it was shown to be an effective dye for the native surface of bacterial spores previously (Xia et al., 2011). The Congo-Red derivative X-34 on the other hand resulted in stains that were clearly distinct between display strains and controls, while the dyes BTD-21 and hFTAA did not show such a clear distinction (see supplemental Fig. 3). Hence, X-34 was chosen for in-depth analysis. 3-D emission/excitation spectra were recorded to determine optimal measurement conditions for the displayed proteins in relation to background generated by the native proteins of the spore-coat. These showed increased fluorescence for TasA-CotY, CotY-SRT10 and CotY-SRT9, when compared to display control strain CotY-SucP (Supplemental Figure 4 B, C, D, E). Results from 3-D scans were used to optimize excitation for 2-D spectra to 373 nm. The optimized 2-D spectrum revealed an expected fluorescence increase of controls PY79 (non-producing strain) and CotY-SucP compared to the dye control, indicating a low degree of interaction between X-34 and crystalline, β-sheet rich structures in the spore-coat (Jiang et al., 2015; Xia et al., 2011). TasA-CotY fluorescence was only marginally increased compared to CotY-SucP. While both TasA-CotY and controls only formed a minor peak at the expected maximum of 444 nm, all SRT-variants showed strongly increased fluorescence. Interestingly, both SRT10-CotY and SRT9-CotY reached higher RFU at 444 nm, than TasA despite no visible protein signal or no visible oligomerization in Western blot analysis respectively. CotY-SRT9 and CotY-SRT10 showed the strongest fluorescence increase and the most distinct peak at 444 nm out of all samples, matching the high degrees of oligomerization observed in Western blot analysis (Fig. 3 A).

**Fig. 3:**
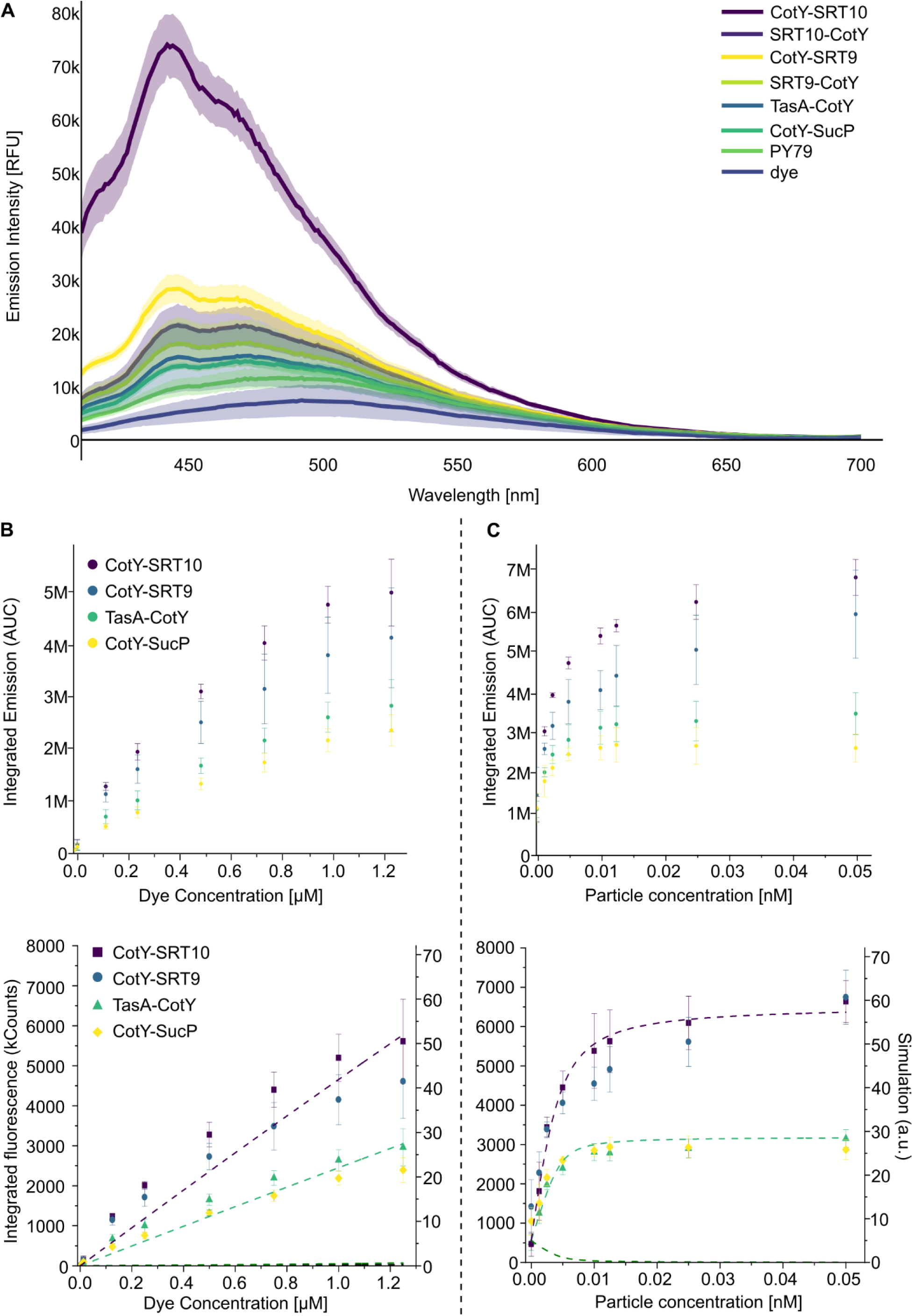
Fluorescence analysis of X-34 dyed spores displaying amyloid proteins. (**A**) 2-D emission spectra obtained for fixed excitation at 373 nm. Y-axis shows emission intensities, and X-axis shows emission wavelengths. Standard deviation is indicated by shading. Averages of 3 biological replicates are shown for all experiments. (**B**) Upper plot: Regression analysis of fixed spore concentration (0.0125 nM) against increasing X-34 concentration indicated on the X-axis. Individual 2-D emission spectra were recorded for each data point. Data points in the graph represent integrals (area under curve, AUC) for each recorded 2-D emission spectrum. Legend shown is relevant for plots B and C. Lower plot: Integrated fluorescence intensity versus X-34 concentration with concentration of particles fixed at 0.0125 nM subjected to Langmuir-type model fit. Dashed lines indicate model fits; markers represent experimental data. (**C**) Upper plot: Regression analysis of increasing spore concentrations indicated on the X-axis against stable X-34 concentration (1.25 µM). Individual 2-D emission spectra were recorded for each data point. Data points in the graph represent integrals (area under curve) for each recorded 2-D emission spectrum. Lower plot: Integrated fluorescence intensity versus concentration of particles with concentration of X-34 fixed at 1.25 mM subjected to Langmuir-type model fit. Dashed lines indicate model fits; markers represent experimental data. Error bars indicate standard deviation.

### 2.2 Protein yield can be estimated using X-34 and mathematical modelling

To approximate the amount of displayed proteins, we conducted a two-way concentration/regression analysis using X-34 (Fig. 3 B, C). First, spores of a concentration of 0.0125 nM were stained using X-34 at rising concentrations from 12.5 nM to 1250 nM. The fluorescence emission spectra of each X-34 concentration were determined, integrated and then plotted. Regression analysis followed the trend of 2-D emission spectra taken at a spore concentration of 0.0125 nM and X-34 concentration of 1250 nM showing a quickly rising slope for CotY-SRT10 and CotY-SRT9. TasA-CotY and CotY-SucP showed a flatter slope at a significantly lower fluorescence level than SRT proteins, although TasA-CotY still increased at a higher rate than control strain CotY-SucP.

In this first regression analysis, slope increase was linear without reaching a plateau for all strains (Fig. 3 B).

The second experiment, a steady dye concentration of 1250 nM was used with spores of increasing particle concentration. This analysis revealed two key findings: First, CotY-SRT10, TasA-CotY and CotY-SucP reached saturation after surpassing particle concentrations of 0.0125 nM, corresponding to the cutoff point of the first experiment. CotY-SRT9 on the other hand did not reach full saturation at this point, instead the slope flattened but continued to increase until the final particle concentration of 0.05 nM. CotY-SRT10, like TasA-CotY did reach full saturation but showed nearly two-fold higher fluorescence intensity than TasA-CotY and CotY-SucP. (Fig. 3 C).

To determine the amount of amyloid-folded protein on the spore-surface data was fitted to a Langmuir-type regression model.

The increased fluorescens of X-34 upon binding to the spore indicates that the binding sites may follow similar binding schemes as pristine amyloid aggregates such as amyloid β (Ikonomovic et al., 2006; Zhang et al., 2018) and transthyretin amyloid fibrils (Sundnes et al., 2025). Amyloid systems can have a multitude of binding sites depending on particular subunits and on symmetry constraints (Todarwal et al., 2021). To test this hypothesis, a rudimentary binding model was introduced assuming *n* equivalent and independent binding sites for each particle class. It was then employed to simulate the results of two fluorescence-based detection assays that varied either the particle or the ligand concentration while holding other sample parameters fixed.

Thus, it is assumed that the spore particles follow the same basic binding kinetics as their pristine amyloid counterparts with primary so called ‘binding sites’ (Ikonomovic et al., 2006; Styren, et al., 2000; Sundnes et al., 2025). Thus, for a particle at concentration [P] with only one binding site and binding to a ligand at concentration [L], the dissociation constant Kd is given by (Bisswanger, 2017):

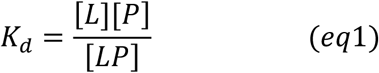

Where *[LP]* is the concentration of ligands *L* bound to *P*. This expression is readily extended to a particle class with *n* equivalent binding sites. Following (Bisswanger, 2017), (details outlined in the SI) this gives the concentration of bound ligands *[LP]* as:

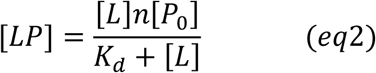

where now *n[P]* represents the total concentration of binding sites. Using mass balance for the total number of ligands [*L*_0_] = [*L*] + [*LP*] eq2 gives a closed expression for the concentration of bound ligands *[LP]* in terms of the total concentration of particles *[P_0_]*, ligands *[L_0_]*, *K_d_* and ‘*n*’; the number of sites per particle in the sample.

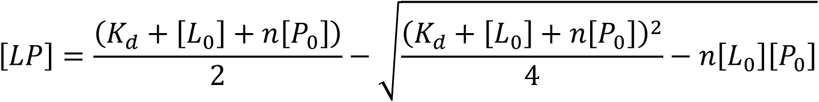

The total fluorescence emission is monitored by either varying *L_0_* or *P_0_*, keeping the other parameter constant (Bisswanger, 2017). For the assessment, the total fluorescence emission is plotted and compared with a simulation adding the calculated values of both *[LP]* and the number of free ligands *[L],* the latter from the mass balance relation. In the simulation model each is contributing with different fluorescence quantum efficiency QE; for L is known (QE_L_ = 4%) from studies of TTR and that of bound ligands *LP* (QE_LP_) is treated as a parameter (Sundnes et al., 2025).

In Figure 3 B and C datapoints are plotted together with simulations using the following parameters; Purple: *K_d_* = 0.13 mM, QE_LP_ = 24%, *n* = 276.000. Light green: *K_d_* = 0.43 mM, QE_LP_ = 43%, *n* = 326.000. It is noted that TasA and SucP both have similar appearances and follow nicely the light green dashed lines, whereas SRT9 and SRT10 both follow closely the purple dashed lines. As judged from the simulated parameters, the latter binds with approx. 3 times lower binding constant (*K_binding_*= 1/ *K_d_*) however, with almost twice the quantum efficiency for bound X34. Both cases have approx. 300.000 binding sites / particle with SRT9 approximately 20% fewer binding sites per particle than SRT10.

**Table 1:** Approximated binding sites per spore calculated from the Langmuir regression model of datasets displayed in Figure 3.

| Strain | TasA-CotY | CotY-SRT10 | CotY-SRT9 |
| --- | --- | --- | --- |
| Binding sites per spore | 276 000 | 326 000 | 300 000 |

Determined X-34 binding sites were equated to displayed protein in amyloid fold under the assumption that each protein copy offers one binding site for X-34.

Binding sites were converted into moles assuming an average of 1.5*10^11^ spores per L:

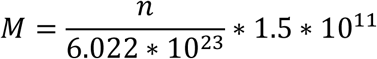

Subsequently molarity was converted into yield (Y) in mg / L using molecular weights of protein monomers:

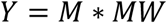

This conversion of model results revealed a general trend towards a yield in the lower mg / L range with the highest yields observed in CotY-SRT10 and TasA (Table 2).

**Table 2:** Approximated yield of amyloid protein calculated from the molarity of protein as determined by X-34 binding sites per spore (see table 1).

| Strain | TasA-CotY | CotY-SRT10 | CotY-SRT9 |
| --- | --- | --- | --- |
| Amyloid yield (mg/L) | 2.2 | 2.2 | 1.6 |

### 2.4 Display of amyloids changes surface structure and stiffness of spores

To further characterize the amyloid proteins displayed on spore surfaces, we employed atomic force microscopy (AFM) to investigate both their surface ultrastructure and nanomechanical properties. The results presented in Figure 4 reveal distinct structural and mechanical signatures associated with different spore types.

**Figure 4:**
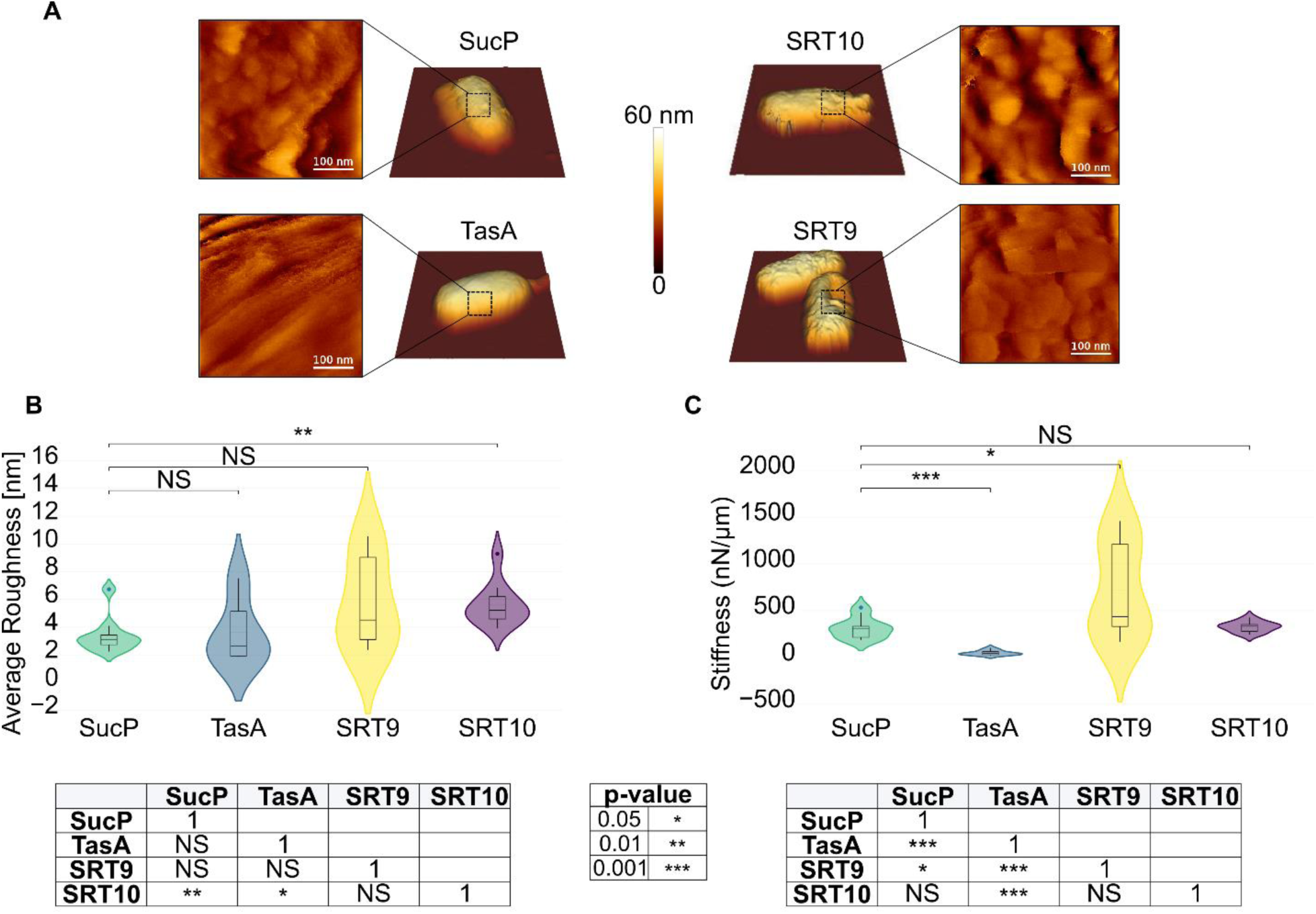
AFM analysis of protein-displaying spores. **(A)** Height images of whole spores displaying the different proteins (central images, color-scale = 1.2 µm). High-resolution images of small areas on top of spores (**400** x **400 n**m²) are shown on the side of each image (color-scales = 60 nm). **(B)** Distribution of roughness values (Ra) extracted from high-resolution images. Measurements in each case were performed on 8-10 spores per spore type. **(C)** Distribution of spring constants values obtained from indentation curves. For each spore type, 8-10 spores were probed. Significance for **(B)** and **(C)** was determined using a Mann-Whitney U test and is indicated according to the resulting p-value. Violin plots with overlaid boxplots display the distribution of roughness values (B) and spring constant values (C).

First, we imaged the spores using Quantitative Imaging™, a force-spectroscopy-based mode that allows high-resolution imaging of samples without damaging them (Chopinet et al., 2013). The images highlighted structural differences among the spores: those presenting SRT proteins appeared rougher than SucP or TasA spores. To quantify these observations, we measured the arithmetic roughness (Ra) across 8–10 spores of each type (Figure 3B). The results confirmed our initial observations, with SRT9 and SRT10 spores exhibiting higher average roughness values of 5.77 ± 3.29 nm and 5.62 ± 1.54 nm, respectively, compared to 3.63 ± 2.33 nm for TasA and 3.38 ± 1.30 nm for SucP. While only the difference between SucP and SRT10 spores reached statistical significance, the greater heterogeneity observed in SRT9 and TasA spores suggests that surface modifications do occur. Notably, SRT-presenting spores displayed globular surface structures, whereas TasA spores exhibited rod-shaped features, further distinguishing their surface morphologies.

Next, we investigated whether the presence of amyloid proteins had an impact on the nanomechanical properties of the spores using nanoindentation experiments. In these experiments, a calibrated AFM tip applies a controlled force to the spore surface, and the resulting deformation is measured to determine its stiffness (spring constant). The results, shown in Figure 3C, revealed that spores possess a rigid envelope, with SucP spores (control) exhibiting an average stiffness of 309.8 ± 120.2 nN/µm, consistent with literature values (e.g., *B. pumilus* spores reported at ∼1000 nN/µm by (Pillet et al., 2016)). However, the expression of different amyloid proteins significantly altered this mechanical property: TasA-presenting spores showed a marked decrease in stiffness (69.0 ± 64.3 N/m), while SRT9 spores exhibited a significant increase (613.8 ± 475.9 N/m). SRT10 spores, meanwhile, retained a stiffness value (326.4 ± 68.8 N/m) comparable to the control. The lower stiffness of TasA spores may be linked to their unique rod-shaped surface structures, which differ from the globular patterns observed in other spores.

### 2.5 Amyloid displaying spores can be used as 3-D printing additive

For 3-D printing spores were dispersed in UV-printing resin at concentrations of 1.5*10^14^ spores / L. Subsequently spore / resin mixtures were printed using a stereolithography 3-D printer into DIN EN ISO 527-2 conform test pieces and post-processed by curing under UV-light at 60 °C. As a reference, samples were printed from pure resin, without the addition of spores.

The test pieces were subjected to maximum tensile strength testing to determine breaking point and elasticity. While adding spores of control strain CotY-SucP increased sample variance, the spores had no significant impact on material tensile strength. TasA-CotY spores increased tensile strength significantly from 51.68 MPa to 56.31 MPa. Adding SRT displaying spores, however, significantly decreased tensile strength to 47.41 and 42.37 MPa for SRT9 and SRT10 respectively. This clearly demonstrates the impact of the displayed amyloid proteins on the printed material.

**Figure 5:**
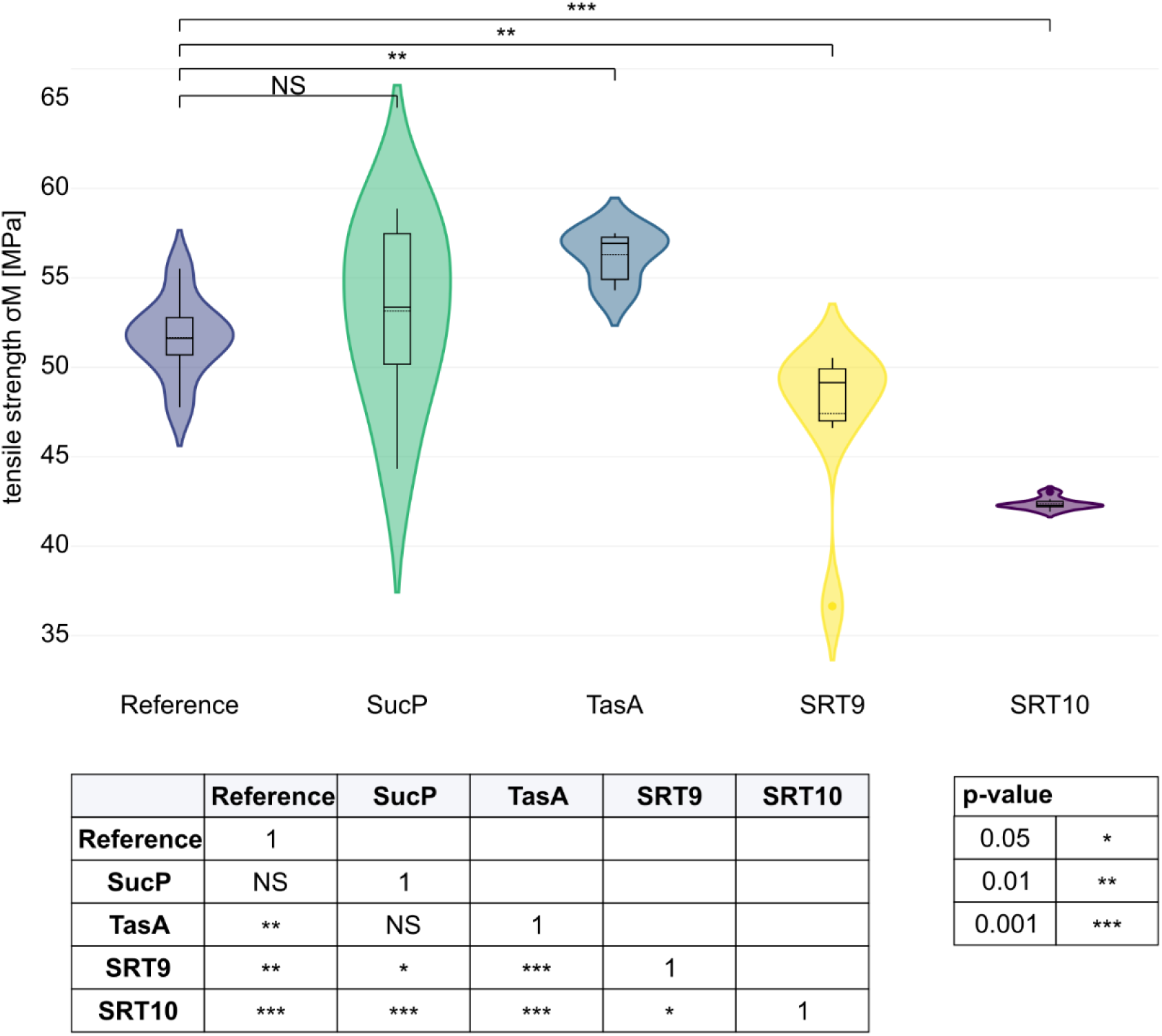
Tensile strength of UV-curable resin composites containing different protein-displaying spores as additives that were dispersed in the resin prior to 3-D printing. After printing, specimens were post-cured under UV-light at 60 °C and subsequently tested in uniaxial tension, with 9 replicates for the unmodified reference resin and 8 replicates for each spore-containing formulation. Violin plots with overlaid boxplots display the distribution of tensile strength (MPa) for the reference, SucP-, TasA-, SRT9-, and SRT10-modified composites, indicating both the spread of the data and the median response for each condition. Significance was determined using a Mann-Whitney U test and is indicated according to the resulting p-value.

## Discussion

In this study, we successfully demonstrate the use of bacterial spores as a production platform for amyloid-like proteins. We could confirm the presence of displayed TasA-, SRT10- and SRT9-variants on the spore-surface via Western blot analysis. Interestingly, no signal was obtained for the C-terminal fusions SRT10-CotY and SRT9-CotY only showed a signal for the protein monomer, while the N-terminal fusions CotY-SRT10 and CotY-SRT9 showed strong overall signals and a high degree of oligomerization. Furthermore, TasA display was only successful for the C-terminal fusion protein (TasA-CotY) and in the presence of auxiliary proteins TapA and SipW. This is not unexpected, since effects on protein display and fusion protein production caused by positioning of fusion protein or purification tags is a well described phenomenon (Structural Genomics Consortium, 2008). Furthermore, TapA has been shown to be essential for TasA oligomerization (Romero et al., 2011). These results confirm that both TasA and SRTs are present on the spore surface, and that amyloid-like proteins produced by spore-surface display can form oligomeric forms that are stable in denaturing conditions. The observation of oligomeric forms on a denaturing SDS-PAGE is not unexpected, as oligomeric amyloid forms of e.g. amyloid-β and Tau from tissue samples have been observed in Western blots prior to this study (Pedrero-Prieto et al., 2019; Scholtzova et al., 2014). However, due to the harsh treatment required for spore-coat stripping, the obtained bands do not give an accurate indication of the native state of the proteins on the spore surface.

Quantitative analysis using fluorescent amyloid probes, especially X-34, revealed that SRT10 when N-terminally fused to CotY (CotY-SRT10) is highly fibrillar, while CotY-SRT9 is at least partially fibrillar. Spectral shift and fluorescence increase were generally stronger in SRT-variants than in TasA-CotY. CotY-SRT10 showed a roughly 2.5-fold increase in fluorescence compared to CotY-SRT9. This is in line with reported protein structures of SRT10 which is partially crystalline with a high amount of cross linked β-barrels while its counterpart SRT9 is reported to be more amorphous and richer in α-helices and random coils (Cámara-Almirón et al., 2023; Guerette et al., 2014). In summary, proteins characterized by a higher degree of structural order and a tendency to form cross-linked β-barrels produce a significantly stronger fluorescence signal upon staining. This was further confirmed by mathematical modeling. While the approach we took is well established for determination of binding sites in single proteins (Sundnes et al., 2025), this is the first time it has been applied to a particle, in this case a spore. It should be noted that this simplified model does not account for differences in binding sites and unspecific binding but gives a rough estimate on the amyloid binding capability. It is noteworthy that values for the binding sites in TTR-amyloid following a similar binding curve analysis were found to have similar values of the quantum efficiency for X-34, however with an approximately 3 – 10 times lower affinity (Sundnes et al., 2025). Yield of actively folded amyloid for all strains was determined to be in the low mg / L range, which is comparable to protein yields for complex proteins such as monoclonal antibodies in other platforms (Zhang et al., 2025). Furthermore, our results strongly suggest that overall protein yield scales with spore density. Hence, yield optimization could be improved through improved fermentation conditions without further engineering of protein production pathways. This method of screening and modeling should be applicable to a wide range of proteins in amyloid fold on a particle surface and offers a novel and rapid way of quantifying displayed protein.

AFM imaging analyses revealed how the expression of amyloid proteins alters the surface architecture of bacterial spores. Spores presenting SRT proteins exhibited globular structures and increased surface roughness, while TasA spores displayed rod-shaped features, marking a clear distinction in protein-specific morphologies. Nanomechanical analysis further demonstrated that the spore surface exhibits rigid behavior, with minimal elastic deformation (<50 nm), precluding the use of the Hertz model (Arnoldi et al., 2000). Instead, a linear force-indentation model was employed to quantify mechanical properties. While TasA expression significantly reduced spore stiffness, SRT9 spores showed a marked increase in rigidity, consistent with the known mechanical properties of these proteins: SRTs behave as rigid, plastic-like materials, whereas TasA forms more flexible fibers (Ghrayeb et al., 2021; Rieu et al., 2016). These findings link protein expression to both structural and mechanical modifications of spore surfaces. SRT proteins reinforce surface rigidity and roughness, while TasA introduces distinct rod-shaped structures that can be correlated with reduced mechanical resilience. The pronounced heterogeneity observed in SRT9 spores suggests a more complex surface organization, further emphasizing the protein-specific impact on spore morphology. These experiments provide the first quantitative validation of how amyloid proteins physically integrate into and modify spore surfaces. Beyond confirming their successful display, these results provide the first biophysical characterization of how amyloid protein expression reshapes spore surfaces, revealing protein-specific modifications in nanomechanical rigidity and surface architecture. These insights could, for example, inform the design and optimization of industrial-scale spore production processes, where surface properties can have an impact on culture stability or downstream processing efficiency.

The use of bacterial spores as material additives and 3-D printing has been demonstrated before. However, to the best of our knowledge, never with a focus on displayed proteins. Instead, the focus was mostly on using spores to survive the harsh conditions of material formation and use the later germinating, living cells as functional element (González et al., 2020; Sidhu et al., 2023; Wüst et al., 2022). In the demonstrated stereolithography 3-D printing process, we could show that the display of amyloid proteins significantly alters the tensile strength of printed test-pieces, while adding control spores had minimal impact on the medium strength but significantly increased the fluctuation of results, indicating that the control spores acted as interfering particles in the resin matrix and had no effect on the structural composition of the printed material. Interestingly, heat treatment during 3-D printing had no significant effect on CotY-SRT10 and CotY-SRT9 (see supplemental Fig. 5), which was unexpected, since SRT proteins have been described to possess thermoplastic properties (Hiew et al., 2016). However, the temperature for breakdown of supramolecular SRT networks has been reported to be as high as 250 °C, a temperature that was not attempted due to the limits of spore and resin stability (Latza et al., 2015). TasA-CotY supplemented resin on the other hand showed significantly increased tensile strength when heat treatment occurred during post curing of the material after printing, hinting at a confirmation change from non-amyloid to amyloid during the post curing process. To our knowledge, TasA aggregation by heat has not been demonstrated before. However, it is well described that TasA can form both amyloidic and non-amyloidic fibrils depending on treatment and conditions (Erskine et al., 2018; Ghrayeb et al., 2021). Furthermore, the use of heat to induce amyloid formation is a well proven method (Lara et al., 2012; Sasahara et al., 2007).

Looking towards the future, we envision two possible future applications of the described system. First as a rapid production platform which circumvents the drawbacks of classical cytosolic production, cell-lysis or secretion. Protein purification could be achieved by a cycle of sporulation and germination with a purification step after germination and degradation of the remaining spore-coat or cleavage from the surface which has been demonstrated before (Spannenkrebs et al., 2025). The obtained protein could then be processed for the generation of novel biomaterials, and the remaining spores recycled as starting biomass for another production round. The other possibility would be to build on the 3-D printing experiments conducted in this study and use the spores themselves as building blocks for biomaterials by forming supramolecular networks with integrated spores which could display additional proteins such as enzymes. This could either be achieved directly by modulation of described amyloid treatment protocols like e.g. wet-spinning and electrospinning, or via the formation of composite materials including amyloid displaying spores, as already demonstrated in this study via stereolithography 3-D printing (Belbéoch et al., 2021; Ki et al., 2007). Another option would be the formation of protein hydrogels derived from e.g. collagen or gelatin in which protein-protein interaction with the displayed amyloids could occur. Research on protein hydrogels and amyloids has thus far been confined to the medical field. However, propagation of amyloids through such hydrogels has been demonstrated (Dalpadado et al., 2016; Simpson et al., 2020). Another interesting candidate molecule for the formation of amyloid-spore hydrogels would be chitosan. A formulation of squid ring teeth proteins and chitosan has been demonstrated before (Chen et al., 2025). This interaction might be of great interest, since chitosan has not only been used in conjunction with bacterial spores before but also shown to inhibit their ability to germinate (Mellegård et al., 2011). Materials produced in such a way could be functionalized with additionally displayed proteins for catalytic applications such as flow-chemistry or even in the medical field.

## 4. Concluding remarks

This study demonstrates bacterial spores as an amyloid production platform. The presence of proteins in amyloid fold on the surface that change spore-surface properties provides proof of principle for this new technology. On a broader scale, this technology serves as a novel tool in the existing amyloid production toolkit. Compared to other amyloid production platforms, it has four key advantages: 1. Simple production and purification by starvation of cells and subsequent centrifugation 2. Avoidance of pitfalls like inclusion body formation due to spatiotemporal control of protein synthesis and display on the outside of the spore 3. The possibility for rapid screening of candidate proteins using amyloid dyes by simple incubation with protein displaying spores. 4. The already in place scale-up to kiloton scale through drop-in into existing industrial spore production processes. This unique combination of advantages opens a new avenue for protein characterization. Especially in the production of load bearing proteins that often consist of repetitive sequences, we envision our technology to be a valuable tool. De-novo sequence generation or sequences generated by random scrambling of repeats or sequence elongation are common in this field (Pena-Francesch et al., 2018; Schmuck et al., 2024; Tokareva et al., 2013). While large sets of sequences can be generated easily, screening of candidate proteins is time-consuming and costly. Our technology could serve as a novel first step in this process, enabling rapid screening of candidate proteins before more rigorous and time-intensive methods of production and testing are used. Moreover, this technology might allow for the production of programmable biomaterials in the future. Materials based on amyloid displaying spores could be fine-tuned by the type of amyloids displayed as demonstrated by the impact of different amyloid-like proteins on stereolithography 3-D printed materials and the change in surface structure and stiffness observed using AFM. Furthermore, materials could be functionalized by co-display of other proteins, such as enzymes.

## 5. Outstanding Questions

1. Is downstream processing in the sense of removing the protein from the spore-surface, purifying it and then forming e.g. fibers feasible?
2. What is the influence of the spore in amyloid based biomaterials?
3. Are composite materials of e.g. protein hydrogels and amyloid displaying spores a mechanically interesting alternative to existing biomaterials?
4. If materials incorporating spores directly are a feasible option: How to deal with the GMO-status of the amyloid displaying spores?

## 6. Materials and Methods

### 5.1 Plasmid generation

Plasmids used in this study were either obtained using Gibson (Gibson et al., 2009) or SLiCE cloning (Zhang et al., 2014) or ordered directly from TWIST bioscience as synthesis DNA. All vectors were based on the replicative high-copy, spore display expression vector pCascade (Karava et al., 2021). Used oligonucleotides are described in supplemental table 1 and vectors are described in supplemental table 2 and are provided as genebank files via the zenodo repository. Enzymes used for cloning were purchased from NEB (Ipswich, MA, USA).

Initial expression vectors for all investigated proteins (p35003, p35004, p98013-98016) were based on insertion of codon-optimized genes encoding for amyloidic target proteins into pCascade as either C- or N-terminal fusions to *cotY in silico* and synthesized by TWIST bioscience (TWIST Bioscience, South San Francisco, CA, USA). Proteins of interest were genetically fused to the spore coat protein CotY which served as an anchor and facilitated localization in the spore crust. Spatiotemporal control of gene expression was regulated using the promoter *P_cotYZ_* which is controlled by sigma^K^ and thus active in the mother cell during the late stage of sporulation (Zhang et al., 1994). CotY and the target proteins were separated by a flexible linker consisting of two glycine-glycine-glycine-glycine-serine repeats (*2xgggs)* and a TEV-protease cleavage site. For Western blot detection, a sequence encoding for an HA-tag was added between the target protein and the flexible linker using Gibson cloning resulting in the generation of p53004 (Field et al., 1988). All vectors were transformed into *E. coli* grown at 37 °C in Lysogeny broth (LB) for maintenance and analysis. Correct sequences were confirmed by colony PCR and whole-plasmid-sequencing (Eurofins Genomics, Ebersberg, Germany). Used oligonucleotides are documented in supplemental table 1; a comprehensive list of all vectors used can be found in supplemental table 2.

### 5.2 Strain generation

Germination deficient strain Bs02003 was used as a progenitor strain for all subsequent strains. High-copy shuttle vectors based on pCascade were transformed into *B. subtilis* using natural competence and rolling circle amplification as previously described (Karava et al., 2021). Rolling circle amplification was carried out using a GenomiPhi V2 DNA amplification Kit (Cytiva, Wilmington, DE, USA). Selection was carried out on LB plates with Neomycin (50 µg / ml). Neomycin of the same concentration was subsequently added to cultures of bacteria harboring shuttle vectors. Genomic integration and deletion were achieved via homologous recombination. Deletion of the *tasA-*operon was facilitated by a vector harboring a Spectinomycin cassette flanked by recognition sites for the Cre-recombinase system (*lox*-sites) and flanking regions homologous to the outside of the *tasA* operon. *tapA/sipW* overexpression was achieved by genomic integration of a synthetic operon consisting of *tapA* and *sipW* under control of stationary phase promoters *P_ylb_* (Yu et al., 2015) or P*_2_* (Feavers et al., 1988) into the *tasA* locus. Deletion and genomic integration vectors were transformed via natural competence without linearization as described before (Nadler et al., 2019). Selection was carried out on LB plates with Spectinomycin (100 µg / ml). Correct strains were confirmed via one-sided colony PCR for shuttle vectors and two-sided colony PCR for genomic integration or deletion. Spectinomycin resistance was removed after confirmation of correct insertion or deletion using Cre/*lox* recombination (Yan et al., 2008). A comprehensive list of all strains used in this study can be found in supplemental table 3.

### 5.3 Cultivation conditions

Bacteria were cultivated in shake flasks using LB at 37 °C and 225 rpm shaking unless indicated otherwise. For strains harboring shuttle vectors antibiotics were added at previously described concentrations.

### 5.4 Sporulation

Sporulation of *Bacillus subtilis* was carried out in shake flasks using 2 x Schaefer’s glucose M media (16 g/L nutrient broth no. 4 (Sigma-Aldrich, St. Louis, MO, USA), 26.8 mM KCl, 2 mM MgSO_4_). 1 mM Ca(NO_3_)_2_, 0.1 mM MnCl_2_, 0.0001 mM FeSO_4_, 5.55 mM glucose and 100 mM MOPS pH 6.8 were added after autoclaving. Cultures were inoculated to an OD_600_ of 0.1 and incubated at 37 °C for 48 h – 72 h at 225 rpm shaking. Subsequently spores and remaining cells were harvested by centrifugation (3000 x g, 10 min) and remaining cells were lysed using lysozyme (0.1 mg / ml) and washed three times using equal culture volume of PBS (0.137 M NaCl, 0.0027 M KCl, 0.01 M Na_2_HPO_4_, 0.0018 M KH_2_PO_4_) at pH 7.2 before resuspension in 1 ml / 50 ml culture of indicated buffer or solvent.

### 5.5 Stripping of the spore-coat

Freshly harvested spores were adjusted to an OD_600_ of 15 in a volume of 150 µl H_2_O and harvested via centrifugation (5.000 x g, 5 min). Subsequently supernatants were removed, and pellets were solved in 100 µl 4 x LAEMMLI buffer (ThermoFisher scientific, Waltham, MA, USA). Samples were subjected to heat treatment at 95 °C for 10 min before bead-beating using glass beads (0.1 µM circumference) in a Retsch CryoMill MM 400 (Retsch GmbH, Haan, Germany) at 30 Hz for 60 min. Supernatants were cleared by centrifugation at 11.000 x g for 30 sec and either directly subjected to SDS-PAGE or frozen at −20 °C for later analysis.

### 5.6 SDS-PAGE and Western blot analysis

To assay the display of proteins on the spore surface, spore-coats of strains carrying HA-tagged versions of the target protein were stripped as described above. 10 µl of supernatant were loaded onto 10 % TRIS/Glycine gels (Invitrogen, Waltham, MA, USA) and subjected to SDS-PAGE. Proteins were transferred onto PVDF membranes using premade Western blotting kits (Bio-Rad, Hercules CA, USA) and detected using primary anti-HA antibody (mouse; 1:3333, Invitrogen, Waltham, MA, USA) and a secondary anti-mouse IgG-horseradish peroxidase conjugate (1:3333, Promega, Madison, WI, USA). Detection was carried out via HRP-chemiluminescence using Clarity^TM^ substrate (Bio-Rad, Hercules CA, USA) and a ChemiDoc^TM^ XRS+ imaging system (Bio-Rad, Hercules CA, USA).

### 5.7 Dye analysis of spores

X-34 fluorescent dye was purchased from Sigma-Aldrich (Sigma-Aldrich, St. Louis, MO, USA). X-34 stock solution was prepared at 2.5 mM in DMSO. Stock solution was stored at 4 °C for two days before use and used for up to two months (storage at 4 °C). For labeling of spores X-34 working solutions of 0.025 µM, 0.25 µM, 0.5 µM, 1 µM, 1.5 µM, 2.0 µM and 2.5 µM were prepared in water. Spore concentration was adjusted to 0.0125 nM in H_2_O. Spores and X-34 working solutions were mixed 1:1 in a final volume of 100 µl resulting in solutions containing 0.0125 µM, 0.125 µM, 0.25 µM, 0.5 µM, 0.75 µM, 1 µM and 1.25 µM X-34 respectively. Spore OD_600_ and X-34 dilutions were prepared manually while black 96-well flatbottom microplates (Greiner AG, Kremsmünster, Austria) were prepared using an EPmotion 5075 pipetting robot (Eppendorf, Hamburg, Germany). To determine optimal dye concentration endpoint measurements were taken at 373 nm excitation and 442 nm emission using a CLARIOstar plate reader (BMG Labtech, Ortenberg, Germany). Subsequently, 2-D emission spectra from 410 – 700 nm were determined at optimal dye concentration using an excitation wavelength of 373 nm. Lastly, 3-D excitation / emission fingerprints were determined at 320 - 414 nm excitation with fixed emission at 450 nm and 420 – 600 nm emission with 373 nm fixed excitation. Determination of binding coefficient and binding-site-per-spore measurements were carried out by measuring full 2-D spectra of dye dilutions described above against spore suspensions of a concentration of 0.0125 nM. Subsequently spore suspensions of concentrations of = 0.00125, 0.0025, 0.005, 0.01, 0.0125, 0.025 and 0.05 nM were tested against a fixed dye concentration of 2.5 µM. Integrals of spectra were determined using Python 3 and plotted with Plotly. Resulting data was analyzed using a Langmuir-type mathematical model described in the results section. The model was implemented using matlab. The code is available in the Zenodo repository found under data availability.

### 5.8 Atomic force microscopy (AFM) experiments

Prior to AFM experiments, spores were diluted in ultra-pure water, and immobilized on Superfrost^TM^ glass slides (J1800AMNZ, Epredia, Germany) for 30 minutes. After that, the glass slide was rinsed using PBS pH 7.4 to remove unattached spores. Height images of the whole cells were recorded using the Quantitative Imaging mode available on the Nanowizard IV XP AFM (Bruker, USA), with MLCT cantilevers (Bruker, nominal spring constant of 0.01 N.m− 1). Images were recorded with a resolution of 150 × 150 pixels using an applied force comprised between 0.8 and 1.5 nN and a z-length comprised between 1 and 6 μm. In all cases, the cantilever’s spring constants were determined using the thermal noise method prior to imaging (Hutter and Bechhoefer, 1993). To obtain high-resolution height images of the spore surface for roughness analysis, small areas of 0.4 × 0.4 μm on top of spores were further recorded at a resolution of 150 x 150 pixels, using an applied force of 1 nN and a z-length of 1 µm. The height images obtained were then analyzed using the data processing software (Bruker, USA) to determine the arithmetic average roughness (Ra). In each condition, at least 8 different images were recorded on 8 different spores.

The mechanical properties of the spores displaying the different proteins were determined using nanoindentation experiments, following previous work from our team (Pillet et al., 2016). For that, the AFM was used in force spectroscopy mode using an applied force comprised between 0.5 and 12 nN depending on the spores, with RFESPG-75 cantilevers (Bruker, nominal spring constant of 3 N/m). The spring constant (rigidity) of the spores was then determined using the equation k*_cell_*= k(S/1-S) where k is the cantilever spring constant and S the slope of the experimental force-distance curve (Arnoldi et al., 2000). The cantilever’s spring constants were determined by the thermal noise method (Hutter and Bechhoefer, 1993). In each condition, at least 8 different spores were probed.

For both roughness and mechanical properties experiments, to assess the significance of the differences observed between the different conditions tested, non-parametric statistical Mann and Whitney U test was used. The differences were considered significant at p-value <0.05.

### 5.9. 3-D printing

To generate sufficient amounts of spores for 3-D printing, *B. subtilis* cells were cultivated in an airlift fermentation setup. 700 ml of 2 x SG (0.1 M MOPS) media were supplemented with 140 µl Antifoam 204 and inoculated with respective strains to an OD_600_ = 0.1 in 1 L Duran® pressure+ bottles. Airlift was generated using commercially available aeration stones for brewing and set to 1 L / bottle / min. Cultures were heated using a standard water bath set to 37 °C. Exhaust air was filtered through 2 % hypochlorite solution in a standard gas washing bottle. The entire setup was operated under a fume hood to avoid hypochlorite exposure.

Before printing 1.5*10^11^ spores per mL were dispersed in Value UV Resin tough clear (PrimaKreator, Malmö, Sweden) using a T25 Ultra-Turrax (IKA-Werke Gmbh & Co. KG, Staufen, Germany) at 18.000 rpm. Resins containing spores were printed into type 1BA test pieces according to DIN EN ISO 527-2 using a Prusa SL1S masked stereolithography (MSLA) printer (Prusa Research a.s., Prague, Czech Republic). A self-built aluminum-tub served as a vat for the printing process. Printing parameters were set to 15 s of UV exposure for the first ten printing layers and 3.5 s for subsequent layers at a layer thickness of 0.05 mm. After printing, test pieces were washed for 5 min in isopropanol in the Form-Wash wash station (Form Labs, Somerville, Massachusetts, USA). After drying at ambient conditions, they were cured under UV light at 60 °C for 2 x 10 min per side in the Form-Cure curing station (Form Labs, see above) to account for potential effects of heating on the interaction between spore displayed amyloids and UV resin. Test pieces were stored for two weeks before testing.

### 5.10 Analysis of 3-D-printed test pieces

3-D printed specimen were analyzed for their tensile strength by mounting them into a clamping setup in a universal material testing machine ZWICK Z050 (ZwickRoell, Ulm, Germany) according to DIN EN ISO 527-2. Tensile strength was calculated from the measured force divided by the sample’s real measured cross-section.

### 5.11 Data analysis

Statistical analysis was carried out using Python 3.0 and data visualization was achieved via Plotly-based python scripts (Plotly, Montreal, Canada). Programming was carried out using Jupyter notebooks (Project Jupyter) while Perplexity AI (Perplexity AI inc. San Francisco, CA, USA) was used to aid in the generation of code for plotting. Inkscape (Free Software Foundation, Inc. Boston, MA, USA) was used to assemble figures and process Images.

## Technology Readiness: TRL 3-4

We present the display of amyloids on the surface of *Bacillus subtilis* spores as a proof-of-concept platform. We demonstrated that *B. subtilis* spores can be used for the display of fibrillar amyloid proteins and that these proteins can be rapidly analyzed using amyloid dyes. Furthermore, we demonstrated that the displayed amyloids influence the surface structure of the spore. This places this technology on a TRL level of 3-4 according to the NASA technology readiness scale.

The upscaling of this technology is a given, as industry scale production of spores is already established, with BASF SE already producing spores in the thousand tons per annum scale. The two biggest hurdles to propel this technology from a screening platform for amyloid proteins to a production or material platform at large scale are a) the unknown influence of spores on materials and b) the regulatory hurdles of GMO spores in some legislations. Future research will have to focus on the crucial question of how to process the amyloid-spores into a product, for which we foresee two routes: 1. Use spores as programmable bioparticles and incorporate them into (bio)materials via amyloids. 2. Separate the displayed proteins from the spores using the system merely as a production platform.

The property of the spore itself could be an inhibiting factor to the quality of a material incorporating spores compared to a material that is only based on amyloid-like proteins. Protein purification from the spore-coat itself is challenging, as it is resistant to protease treatment and highly resistant to mechanical forces and chemicals, making regular protein purification improbable. Amyloid-like proteins, however, might offer a unique opportunity in this regard, as this class of protein is not as susceptible to structural loss during the harsh treatments required for spore-coat stripping.

## Data availability

All raw data and scripts for their evaluation are provided in a Zenodo repository 10.5281/zenodo.17017715.

## Acknowledgements

We want to thank the Novo Nordisk Foundation for funding this project. C.D., M.P., M.K. and J.K. were funded by the Novo Nordisk Foundation Ascending-Investigator-Grant: “PolySpore - bacterial spores as versatile, programable bulk-biomatter” Grant ID: NNF22OC0073221, acquired by J.K. Additionally we want to thank Professor Catherine Taylor Nordgard for great scientific discussion surrounding the topic of spore-based biomaterials and Jan Benedict Spannenkrebs for help in troubleshooting in the early stages of the project.

## Author contributions

C.D.I., M.P., and M.K. generated strains and plasmids used in this study. C.D.I. and M.P. planned and conducted spore production and characterization. M.L. provided amyloid dyes, advised on their use, and provided key knowledge for quantification. M.L. developed the mathematical model. C.F.D. and S.S. planned and conducted AFM experiments. D.S. and A.U. planned and conducted 3-D printing and material testing. C.D.I. designed and generated figures. M.P. wrote the initial manuscript with input from J.K. All authors contributed to the final manuscript. J.K. and M.P. coordinated the project. J.K. acquired the funding.

