## Supplemental Material for "Bacterial Spores as a Scalable, Modular Platform for the Production of Amyloids for Materials"

Supplemental Table 1: Oligonucleotides used

| Oligo number | Sequence | Use |
|---|---|---|
| o02215 | GAGCTAGTTGACAGAACATCG | Confirmation of Bs98090, Bs98091, Bs98092, Bs98093 |
| o14013 | CGACTCACTATAGGGAGAGCGGC | Confirmation p53002 and p53003 |
| o14127 | CACTGCCCGCTTTCCAG | Confirmation p53002 and p53003 |
| o14159 | GCAGCAGATTACGCGCAG | Confirmation of Bs98090, Bs98091, Bs98092, Bs98093, Bs98106, Bs98107, Bs98108, Bs98109 |
| o29106 | ATAGACGTTCACCCACACCAAGTGGG | Confirmation of p53004 |
| o29153 | AGACAGTGTTGCACAGCAATCG | Confirmation of p53004 |
| o29168 | AAGATGTTGCTGTCTCCCA | Generation of p98035, p98036, p98037 and p98038 |
| o29169 | GAACTTGTCTTTTCCCACG | Generation of p98035, p98036, p98037 and p98038 |
| o53005 | CAATAACCATCGAGACGGCCCAGTAATCACGA ATTGGATCACTTCTCAAAGATCCCATGTGCT | Generation p53002 |
| o53006 | TTGTGAAACAATCGAAACATAC AAATCTCCCCCTTTGTTGT | Generation p53002 |
| o53007 | GCAATGAAGCTGATCAGTAAATG AAGCTGATCAGTAATATTT | Generation p53002 and p53003 |
| o53008 | CGTATAATGTATGCTATACGAACGGTAGGCCTC GAGGATCTTAAGTAGACATGGTGCTGTCCT | Generation p53002 and p53003 |
| o53009 | CAACAAAGGGGGAGATTTGTATGT TTCGATTGTTTCACAATCAGC | Generation p53002 |

| | | |
|---|---|---|
| o53010 | ATATTACTGATCAGCTTCATTT<br>ACTGATCAGCTTCATTGCT | Generation p53002 and p53003 |
| o53011 | CGAGACGGCCCAGTAATCACGAATTG<br>GATCAGGATTTTTTGTGTCATTGGCGA | Generation of p53003 |
| o53012 | AAACAATCGAAACATTTATGT<br>ATCCCTCCATAACGGTTGC | Generation of p53003 |
| o53013 | TGGAGGGATACATAAATGTTT<br>CGATTGTTTCACAATCAGC | Generation of p53003 |
| o53016 | CCCTTACGACGTTCCGGA | Confirmation of Bs98106, Bs98107, Bs98108, Bs98109 |
| o66019 | GGTTGAATTAATGGTGGAAGC | Confirmation Bs53003, Bs53013, and Bs53014 |
| o66020 | AAAGAGGTGCGGAAAGAAAC | Confirmation Bs53003, Bs53013, and Bs53014 |
| o98151 | TACCCTTACGACGTTCCGGATTA<br>TGCAGTTCCACACCTGCTCGCG | Generation of p98035 |
| o98152 | TGCATAATCCGGAACGTCGTAAGGGT<br>ATTGAAAATACAAATTTTCGCTTCCTCC | Generation of p98035, p98037 |
| o98153 | TACCCTTACGACGTTCCGGATTATGC<br>AGAAAATTTGTATTTTCAAGGGGGGG | Generation of p98036, p98038 |
| o98154 | TGCATAATCCGGAACGTCGTAAG<br>GGTAGTGCAGAAAGCCGCCATAC | Generation of p98036 |
| o98155 | TACCCTTACGACGTTCCGGATTATGC<br>ATATCACGCGAACCATGTAGGAAC | Generation of p98037 |
| o98156 | TGCATAATCCGGAACGTCGTAAGGGTAGT<br>TCACCATTTTAATTTCATAGTGTTTACAG | Generation of p98038 |

Supplemental Table 2: Plasmids used in this study

| **Plasmid number** | **Genetic information** | **Method** | **Source** |
|---|---|---|---|
| p02033 | ORI: *colE1* (Ec), *repE* (Bs); KanR*; resolvase beta*;$P_{cotYZ}$,*cotY-(2xgggs)-sucP_bifidobacterum* | SLiCE; shuttle vector | Marianna Karava |
| p35004 | ORI: *colE1 (Ec), repE (Bs);* KanR*; resolvase beta;*$P_{cotYZ}$,*tasA-(2xgggs)-cotY* | Gene synthesis; shuttle vector | This study |
| p98013 | ORI: *colE1 (Ec), repE (Bs);* KanR; *resolvase beta;*$P_{cotYZ}$,*cotY-(2xgggs)-suckerin 10* | Gene synthesis; shuttle vector | This study |
| p98014 | ORI: colE1 (Ec), repE (Bs); KanR; *resolvase beta;*$P_{cotYZ}$*-suckerin10-(2xgggs)-cotY* | Gene synthesis; shuttle vector | This study |
| p98015 | ORI: colE1 (Ec), repE (Bs); KanR; *resolvase* | Gene synthesis; shuttle vector | This study |

| | | | |
|---|---|---|---|
| | *beta;*$P_{cotYZ}$*,cotY-(2xgggs)-suckerin 9* | | |
| p98016 | ORI: colE1 (Ec), repE (Bs); KanR; resolvase beta;$P_{cotYZ}$,*suckerin9-2xgggs)-cotY* | Gene synthesis; shuttle vector | This study |
| p98035 | ORI: *colE1 (Ec), repE (Bs);* KanR; *resolvase beta;*$P_{cotYZ}$*,cotY-TEV-(2xgggs)-HA-suckerin 10* | Gibson cloning; shuttle vector | This study |
| p98036 | ORI: colE1 (Ec), repE (Bs); KanR; *resolvase beta;*$P_{cotYZ}$*-suckerin10-HA-(2xgggs)-TEV-cotY* | Gibson cloning; shuttle vector | This study |
| p98037 | ORI: *colE1 (Ec), repE (Bs);* KanR; *resolvase beta;*$P_{cotYZ}$*,cotY-TEV-(2xgggs)-HA-suckerin 9* | Gibson cloning; shuttle vector | This study |
| p98038 | ORI: *colE1* (Ec), *repE* (Bs); KanR; *resolvase beta;*$P_{cotYZ}$*-suckerin9-HA-(2xgggs)-TEV-cotY* | Gibson cloning; shuttle vector | This study |
| p14076 | ORI: *colE1* (Ec); AmpR; pJET; *sinR ; lox66-spectinomycinR-lox71; yqzG* | integrative | Internal strain collection |
| p53002 | ORI: *pMB1* (Ec); AmpR; *sinR ; pylb::tapA; sipW; ygzG* | Gibson cloning; integrative | This study |
| p53003 | ORI: *pMB1* (Ec); AmpR; *sinR ; p2::tapA; sipW; ygzG* | Gibson cloning; integrative | This study |
| p53004 | ORI: *colE1* (Ec), *repE* (Bs); KanR; *resolvase beta;*$P_{cotYZ}$*-tasA-HA-(2xgggs)-TEV-cotY* | Gibson cloning; shuttle vector | This study |

Supplemental table 3: Strains used in this study

| **Strain number** | **Genotype** | **Abbreviation** | **Strain background** | **Source** |
|---|---|---|---|---|
| PY79 | *Wildtype* | PY79 | - | Bacillus subtilis genetic stock center (BGSC) |
| Bs02003 | *ΔsleBΔcwlD;lox72* | Progenitor strain | Bacillus subtilis KO7 | (Karava et al., 2019) |
| Bs53003 | *ΔtasAΔtapAΔsipW;lox72* | ΔtasA | Bs02003 | This study |
| Bs53005 | Plasmid*:* $P_{cotYZ}$*::tasA-(2xgggs)-TEV-cotY;kanR* | ΔtasA TasA-CotY | Bs53003 | This study |
| Bs53013 | *pylb::tapA;sipW;lox72* | pylb_tapA/sipW | Bs53003 | This study |
| Bs53014 | *p2::tapA;sipW;lox72* | p2_tapA/sipW | Bs53003 | This study |
| Bs53015 | Plasmid*:* $P_{cotYZ}$*::tasA-(2xgggs)-TEV-cotY;kanR* | TasA-CotY-pylb | Bs53013 | This study |
| Bs53016 | Plasmid*:* $P_{cotYZ}$*::tasA-(2xgggs)-TEV-cotY;kanR* | TasA-CotY-P2 | Bs53014 | This study |
| Bs02040 | Plasmid*:* $P_{cotYZ}$*::cotY-(2xgggs)-TEV-sucP;kanR* | CotY-SucP | Bs02003 | Marianna |
| Bs35003 | Plasmid*:* $P_{cotYZ}$*::cotY-(2xgggs)-TEV-tasA;kanR* | CotY-TasA | Bs02003 | This study |

| | | | | |
|---|---|---|---|---|
| Bs35004 | Plasmid*: P$_{cotYZ}$::tasA-(2xgggs)-TEV-cotY;kanR* | TasA-CotY | Bs02003 | This study |
| Bs98090 | Plasmid*: P$_{cotYZ}$::cotY-(2xgggs)-TEV-suc10;kanR* | CotY-Suc10 | Bs53003 | This study |
| Bs98091 | Plasmid*: P$_{cotYZ}$::suc10-(2xgggs)-TEV-cotY;kanR* | Suc10-CotY | Bs53003 | This study |
| Bs98092 | Plasmid*: P$_{cotYZ}$::cotY-(2xgggs)-TEV-suc9;kanR* | CotY-Suc9 | Bs53003 | This study |
| Bs98093 | Plasmid*: P$_{cotYZ}$::suc9-(2xgggs)-TEV-cotY;kanR* | Suc9-CotY | Bs53003 | This study |
| Bs53017 | *Plasmid: P$_{cotYZ}$::TasA-HA-(2xgggs)-TEV-cotY;kanR* | ΔtasA TasA-HA-CotY | Bs53003 | This study |
| Bs53019 | *Plasmid: P$_{cotYZ}$::TasA-HA-(2xgggs)-TEV-cotY;kanR* | TasA-HA-CotY | Bs53014 | This study |
| Bs98106 | Plasmid*: P$_{cotYZ}$::cotY-(2xgggs)-TEV-HA-suc9;kanR* | CotY-HA-Suc9 | Bs02003 | This study |
| Bs98107 | Plasmid*: P$_{cotYZ}$::suc9-HA-(2xgggs)-TEV-cotY;kanR* | Suc9-HA-CotY | Bs02003 | This study |
| Bs98108 | Plasmid*: pcotYZ::cotY-(2xgggs)-TEV-HA-suc10;kanR* | CotY-HA-Suc10 | Bs02003 | This study |
| Bs98109 | Plasmid*: P$_{cotYZ}$::suc10-HA-(2xgggs)-TEV-cotY;kanR* | Suc10-HA-CotY | Bs02003 | This study |

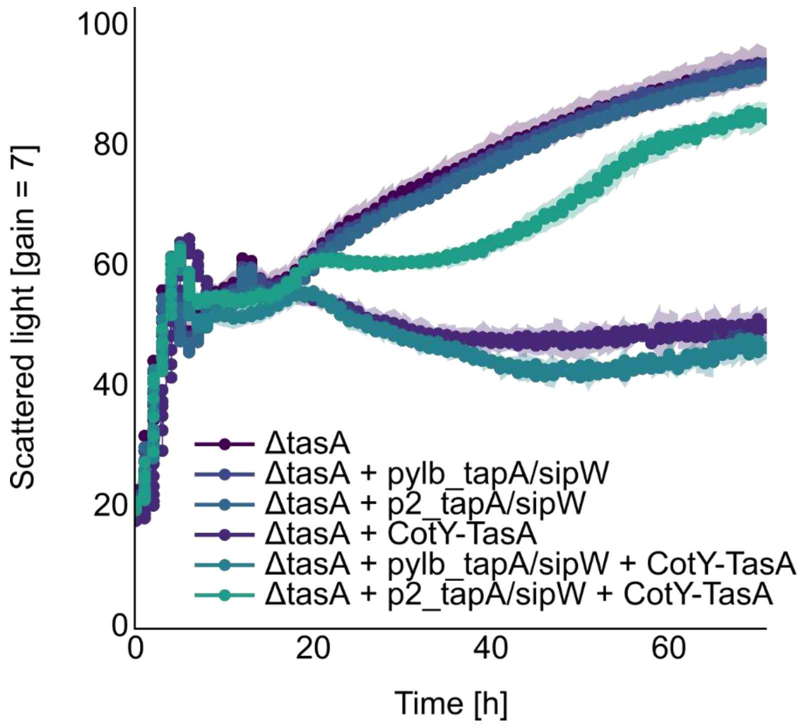


**Supplemental figure 1:** Impact of *tasA* deletion and *tapA* reconstitution on strain fitness. Sequentially generated strains harboring alterations to the *tasA* operon were analyzed in terms of proliferation and sporulation behavior. Cultivations were conducted in 0.1 M MOPS buffered 2 x SG media in Flowerplates™ using a BioLectorPro™ (Beckmann coulter life sciences, Brea, CA, USA). Y-axis indicates scattered light intensity in arbitrary units; X-axis indicates time in hours. Deletion of *tasA* and reintroduction of *tapA* and *sipW* (*pylb_tapA/sipW, p2_tapA/sipW*) had no impact on strain fitness, while strains harboring *tasA* deletion and the *coty-tasA* display plasmid showed strongly reduced overall strain fitness. Reconstitution of high levels of *tapA* and *sipW* using a genomic integration of these genes under control of the strong p2 promoter (high tapA) mostly restored strain fitness in the presence of the *coty-tasA* display plasmid.

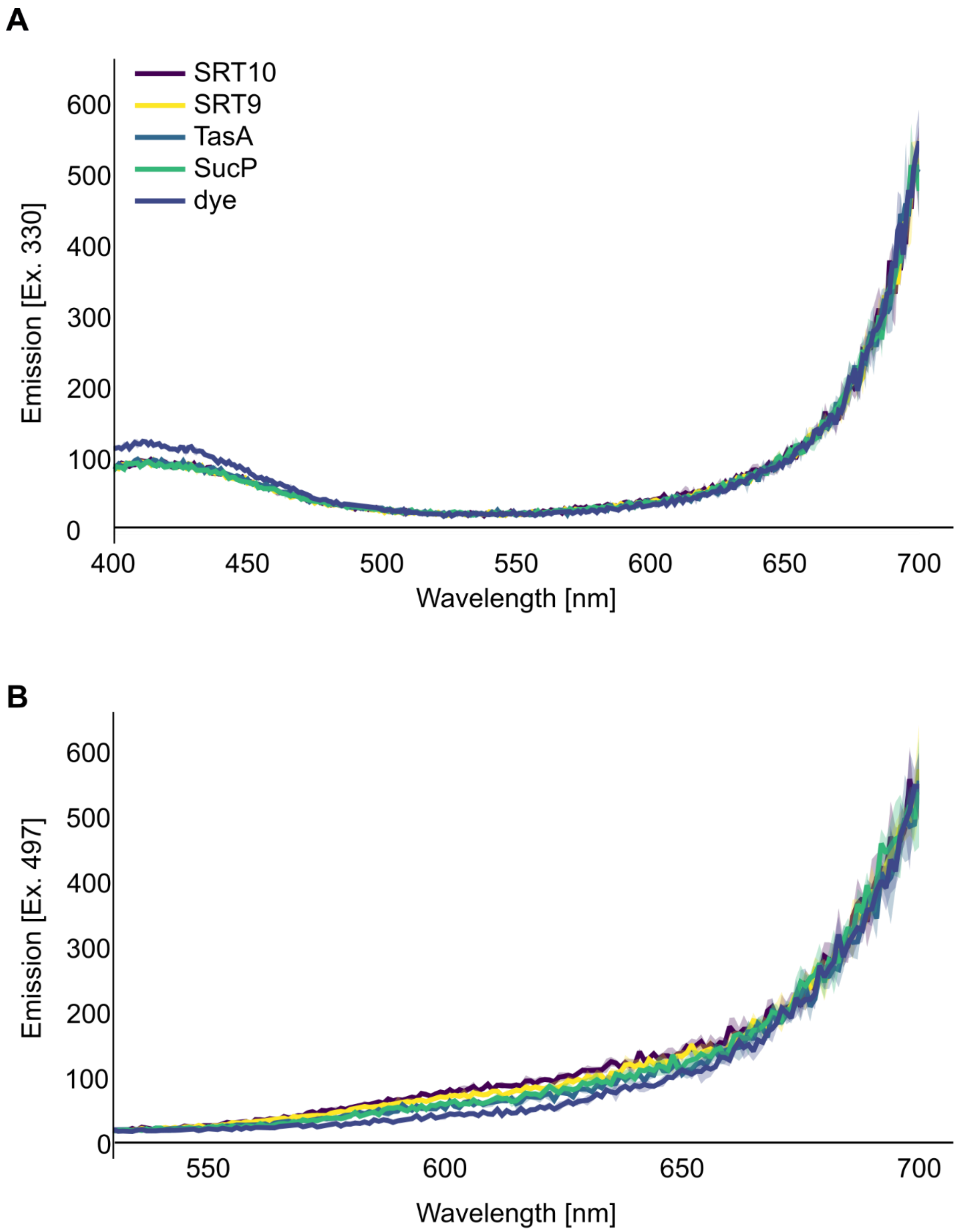


**Supplemental figure 2:** Staining of spores with CongoRed. No distinction could be made between control spores and spores displaying amyloid proteins (**A**) Fluorescence emission spectra of amyloid displaying spores and control (SucP) stained with congored. Emission spectrum was recorded at an excitation of 330 nm, corresponding to unbound CongoRed. All samples showed the same low peak around 420 nm and an increase in fluorescence from 650 nm onward, but no distinction between samples could be made. (**B**) Fluorescence emission spectra of amyloid displaying spores and control (SucP) stained with CongoRed. Emission spectrum was recorded at an excitation of 497 nm, corresponding to bound CongoRed. No peak could be detected but an increase in fluorescence from

650 nm onward was observed for all samples. However, no distinction between samples could be made.

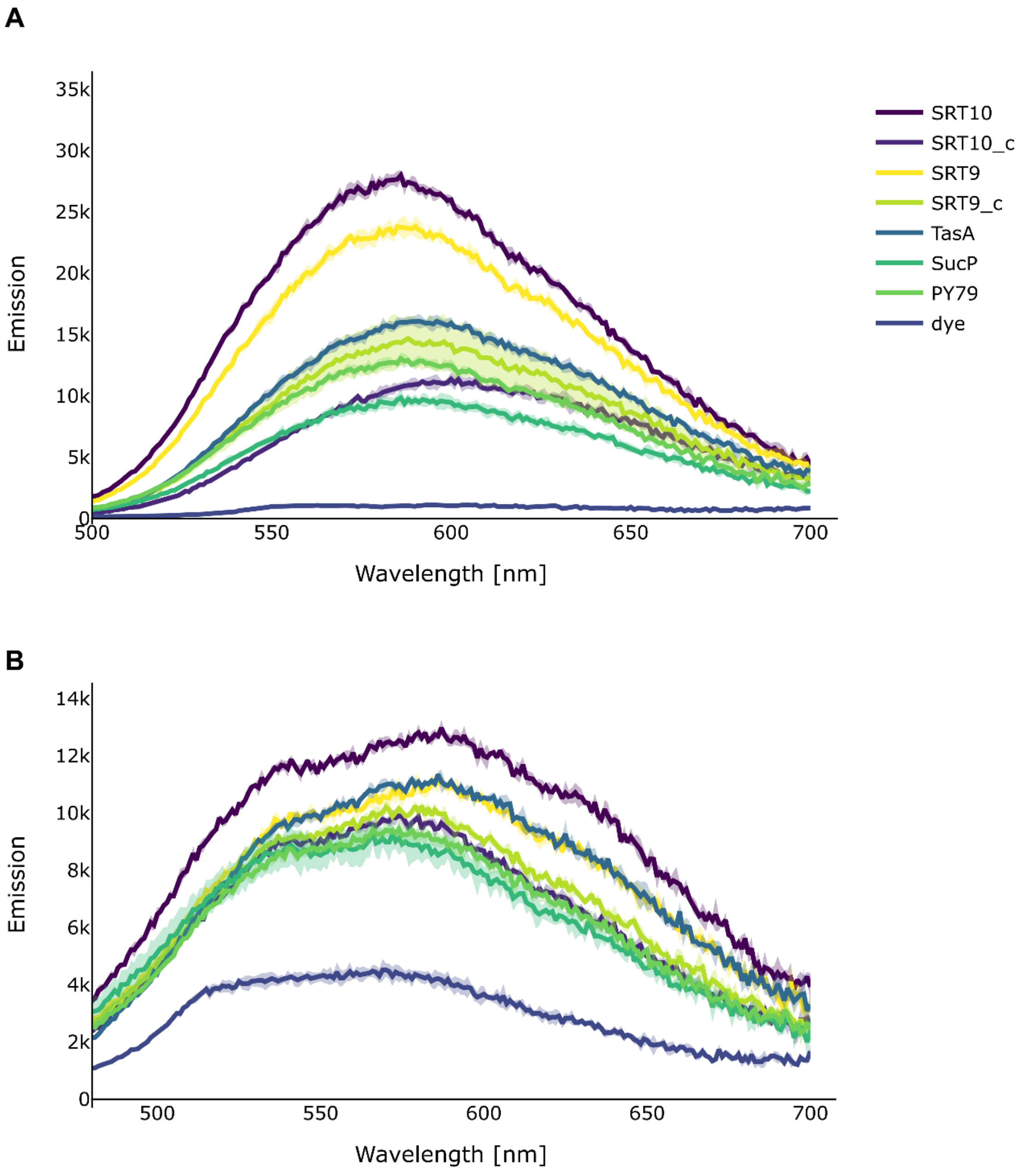


**Supplemental figure 3:** Staining of spores with amyloid specific dyes BTD-21 (A) and hFTAA (B). Staining with both dyes followed a similar trend to staining with X-34 although less pronounced, hence X-34 was chosen for more in-depth analysis. (**A**) Fluorescence emission spectra of amyloid displaying spores and controls (PY79 and SucP) stained with fluorescent dye BTD-21. Overall amyloid displaying strains showed stronger signals than control strains. However, no clear shift in wavelength was observable, since all samples showed a maximum around 595 nm. Other than X-34 staining (Fig. 2) and Western bot analysis (Fig. 1) TasA displaying spores showed a signal almost as strong as that of spores displaying SRT10. (**B**) Fluorescence emission spectra of amyloid displaying spores and controls (PY79 and SucP) stained with fluorescent dye hFTAA. Fluorescence emission spectra follow the same trend as Western blot analysis and X-34 staining with CotY-SRT10 and CotY-SRT9 displaying spores showing the highest fluorescence peaks. Like staining with hFTAA no shift in wavelength was observed between samples and controls.

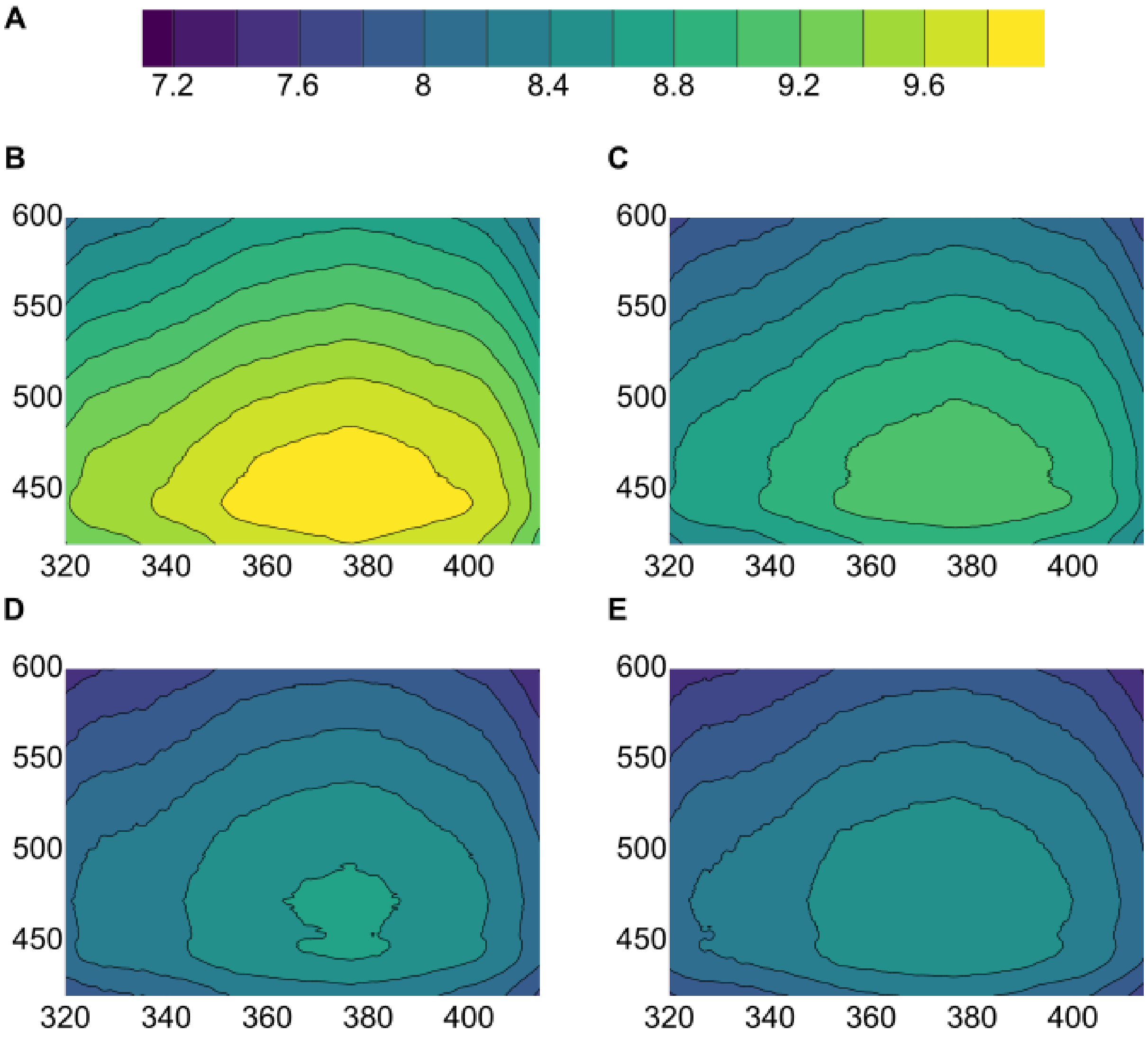


**Supplemental figure 4:** Fluorescence analysis of X-34 dyed spores displaying amyloid-like proteins (**A**) Color scale indicating logarithmic emission intensity in 3-D fingerprint plots. (**B-E**) 3-D fingerprint plots obtained from Emission/Excitation scans. Excitation wavelength shown on X-axis, emission wavelength shown on Y-axis. Emission intensity visualized through color scale described in figure 3 A. (**B**) X-34 dyed spores of strain displaying CotY-SRT10. (**C**) X-34 dyed spores of strain displaying CotY-SRT9. (**D**) X-34 dyed spores of strain displaying TasA-CotY. (**E**) X-34 dyed spores strain displaying negative control protein CotY-SucP. Analysis revealed a shared excitation maximum at 373 nm for all strains, while fingerprint shape and recorded fluorescence intensity were distinctly different.

**A**

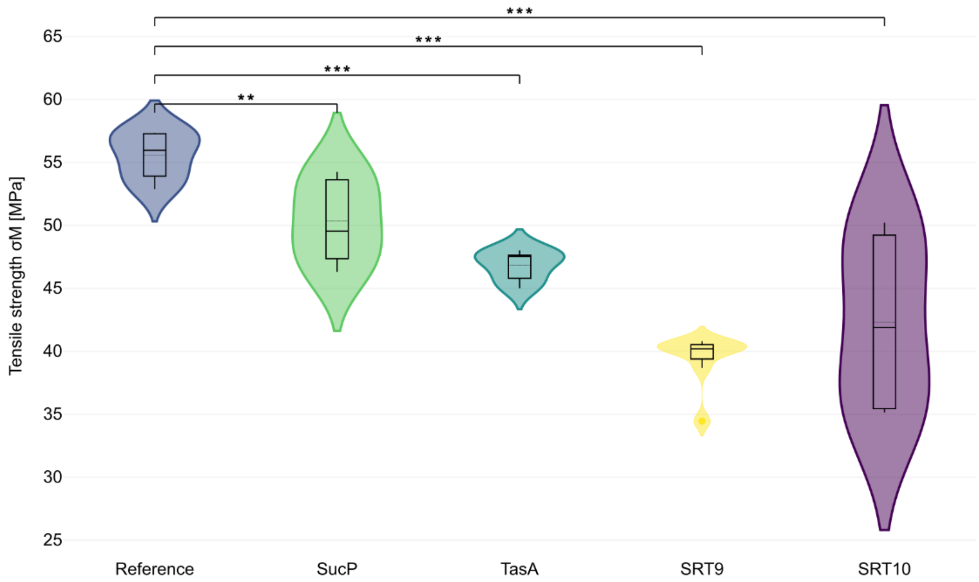


| | Reference | SucP | TasA | SRT9 | SRT10 |
|---|---|---|---|---|---|
| **Reference** | 1 | | | | |
| **SucP** | ** | 1 | | | |
| **TasA** | *** | NS | 1 | | |
| **SRT9** | *** | *** | *** | 1 | |
| **SRT10** | *** | * | NS | NS | 1 |

**B**

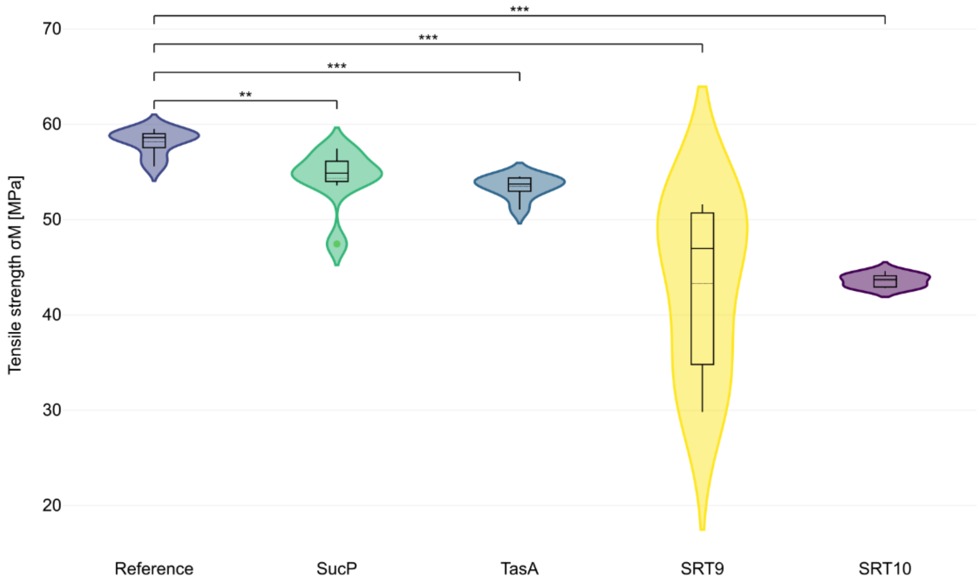


| | Reference | SucP | TasA | SRT9 | SRT10 |
|---|---|---|---|---|---|
| **Reference** | 1 | | | | |
| **SucP** | ** | 1 | | | |
| **TasA** | *** | NS | 1 | | |
| **SRT9** | *** | ** | *** | 1 | |
| **SRT10** | *** | *** | *** | NS | 1 |

**Supplemental figure 5:** Tensile strength of UV-curable resin composites containing different protein-displaying spores as additives that were dispersed in the resin prior to 3-D printing. For specimens in **(A)** the UV-curable resin composites were heated to 60 °C before printing and printed specimen were post-cured after printing under UV-light at RT, while for specimens in **(B)** the UV-curable resin composites were heated to 60 °C before printing as well, but the printed specimen were post-cured under UV-light at 60 °C. The samples were then tested in uniaxial tension, with 9 replicates for the unmodified reference resin and 8 replicates for each spore-containing formulation. Violin plots with overlaid boxplots display the distribution of tensile strength (MPa) for the reference, SucP-, TasA-, SRT9-, and SRT10-modified composites, indicating both the spread of the data and the median response for each condition. Significance was determined using a Mann-Whitney U test and is indicated according to the resulting p-value.

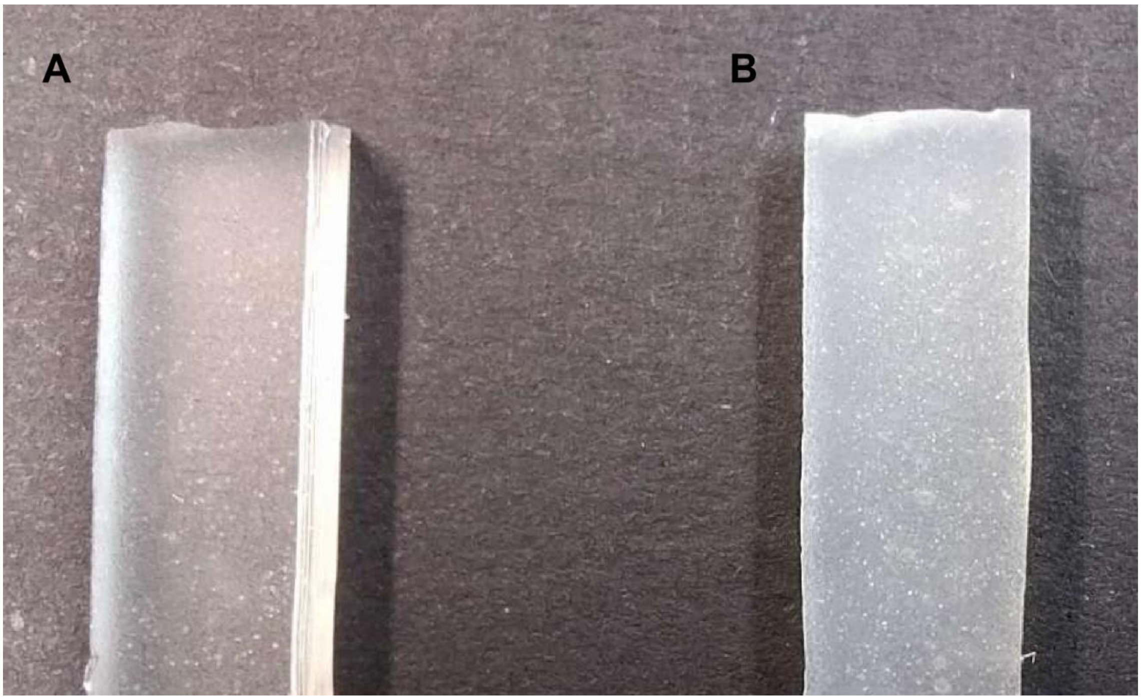


**Supplemental figure 6:** Photographs of 3D-printed test pieces prior to tensile testing. (**A)** Control specimen fabricated using native resin. (**B)** Functionalized composite specimen containing resin infused with amyloid-like protein displaying spores. Specimens were printed according to standard geometries for mechanical characterization.
